# The large majority of intergenic sites in bacteria are selectively constrained, even when known regulatory elements are excluded

**DOI:** 10.1101/069708

**Authors:** Harry A. Thorpe, Sion Bayliss, Laurence D. Hurst, Edward J. Feil

## Abstract

There are currently no broad estimates of the overall strength and direction of selection operating on intergenic variation in bacteria. Here we address this using large whole genome sequence datasets representing six diverse bacterial species; *Escherichia coli*, *Staphylococcus aureus, Salmonella enterica*, *Streptococcus pneumoniae*, *Klebsiella pneumoniae*, and *Mycobacterium tuberculosis*. Excluding *M. tuberculosis*, we find that a high proportion (62%-79%; mean 70%) of intergenic sites are selectively constrained, relative to synonymous sites. Non-coding RNAs tend to be under stronger selective constraint than promoters, which in turn are typically more constrained than rho-independent terminators. Even when these regulatory elements are excluded, the mean proportion of constrained intergenic sites only falls to 69%; thus our current understanding of the functionality of intergenic regions (IGRs) in bacteria is severely limited. Consistent with a role for positive as well as negative selection on intergenic sites, we present evidence for strong positive selection in *Mycobacterium tuberculosis* promoters, underlining the key role of regulatory changes as an adaptive mechanism in this highly monomorphic pathogen.

## Introduction

The ability to generate whole genome sequence datasets from very large samples of bacterial isolates recovered from natural populations provides unprecedented power to dissect evolutionary processes. Although tests for selection are routinely carried out on the ~85-90% of bacterial genomes corresponding to protein-coding genes, the strength and direction of selection operating on intergenic regions (IGRs) tends to be overlooked ^1^. Indeed, in many cases the presumption is that intergenic sites provide a valid proxy for neutrality ^2–4^. Although some studies have investigated selection on intergenic sites in bacteria ^1,5,6^ these are relatively scarce, and the perception that mutational changes in IGRs are generally neutral and have little or no selective cost still widely persists ^3,4^. This is clearly evidenced by the recent establishment of bacterial whole genome database infrastructure that excludes, by default, IGRs altogether ^7–11^. An exception is the work by Fu *et al,* who generated a core genome consisting of genes and IGRs for *Salmonella enterica* serovar *Typhimurium*, and demonstrated that the inclusion of IGRs in the core genome increased discriminatory power to differentiate outbreak isolates ^4^.

The standard approach to measuring selection, the ratio of non-synonymous to synonymous changes (dN/dS), is clearly not valid for IGRs, and the perceived lack of established methodology can in part explain the rather casual dismissal of the adaptive relevance of intergenic variation. Can we continue to maintain the presumption of neutrality of IGRs? The paucity of studies aimed at systematically addressing this question is strikingly at odds with the many recent examples demonstrating the phenotypic impact of mutations in riboswitches, small RNAs, promoters, terminators, and regulator binding sites ^12^. Single nucleotide polymorphisms (SNPs) or small insertion/deletions (INDELs) within these elements can have major phenotypic consequences. For example, in a recent GWAS study, 13 intergenic SNPs were found to be significantly associated with toxicity in Methicillin-resistant *Staphylococcus aureus* (MRSA), and four of these were experimentally validated ^13^. In *Mycobacterium tuberculosis*, mutations within the *eis* promoter region increase expression of Eis, an enzyme which confers resistance to kanamycin and promotes intracellular survival ^14^.

In addition to those studies focussing on naturally occurring mutations, knock-out experiments on regulatory RNAs have also confirmed their key roles in virulence and other important phenotypes such as competence. For example, the *Salmonella* sRNA IsrM is important for invasion of epithelial cells and replication inside macrophages ^15^. In *S. aureus*, the Sigma B-dependent *RsaA* sRNA represses the global regulator *MgrA*; this decreases the severity of acute infection and promotes chronic infection. ^16^ In *S. pneumoniae*, the *srn206* non-coding RNA is involved in competence modulation ^17^.

These well characterised regulatory elements are clearly expected to be under strong selective constraint, but to what extent are these examples typical with respect to IGRs in general? If such functional sites are rare a presumption that the great bulk of IGR sequence is neutrally evolving and functionless is sustainable. Molina *et al* analysed 22 diverse bacterial clades and found evidence for purifying selection on non-coding regions of the genome, the strength of which was related to the presence of transcription factor binding sites ^1^. Luo *et al* analysed 13 group A streptococcus genomes, and found evidence for ongoing purifying selection on intergenic sites ^5^. Similarly, a study by Degnan *et al* found considerable conservation of intergenic sequences in *Buchnera* genomes ^6^. *Buchnera* are obligate symbionts and display typical features of genome reduction such as high genomic AT content, gene loss, and pseudogenisation. These genomic properties reflect inefficient selection (or equivalently a high rate of genetic drift) resulting from intracellular lifestyles and small effective population sizes. Neutral intergenic sites in endosymbiont genomes are therefore expected to be rapidly degraded and deleted; the fact that intergenic sequences appear to have been maintained in *Buchnera* is therefore strong evidence that they are indeed functional.

Despite these studies, there remain no broad quantitative estimates of the commonality of selective constraint operating on IGRs, which intergenic regulatory elements are under strongest selection, nor whether a signal of selection can be detected even for those intergenic sites for which there is no known function. Here we use two independent approaches to address these questions, one based on the established logic of site frequency spectra (the Proportion of Singleton Mutations; PSM), and the other a modification of dN/dS (dI/dS; where dI is the number of intergenic SNPs per intergenic site).

According to the classical nearly neutral theory, deleterious mutations are not always eliminated immediately from a population, but can persist for a period of time determined by the selection coefficient and the effective population size ^18,19^. If mutations are only very weakly deleterious they might increase in frequency by drift, but be purged after a considerable lag, whereas more severe deleterious mutations will be lost more quickly whilst they are still very rare. The rarest SNPs are those that are observed in only one genome (singletons), thus the proportion of singleton mutations (PSM) can be used to gauge the frequency of highly deleterious mutations. Note that the most deleterious mutations of all are those that are lethal and hence never observed. Excepting those cases where the effective population size is very small, the weaker effect mutations will tend to be lost over time, and a reduction in dN/dS (or, equivalently, dI/dS) with increasing divergence acts as a footprint of weak purifying selection ^20^. Consequently, dN/dS and dI/dS from sufficiently diverged genomes make it possible to estimate the proportion of non-synonymous or intergenic mutations that have been selectively purged, relative to synonymous sites. This will likely be a conservative estimate for the proportion of sites under selective constraint for two reasons. First, it is unlikely that the comparator (dS) is a perfect neutral benchmark, synonymous mutations being under, for example, translational selection (operating on codon usage bias ^21^). Second, dI/dS and dN/dS will continue to decrease with divergence time but potentially rise as saturation of synonymous sites becomes more evident. As there is no reason to suppose that the available data will correspond to the minima for a given species, we will tend to over-estimate dI/dS and dN/dS. Whereas the PSM approach provides a measure of how many SNPs are purged very quickly due to highly deleterious effects, measures of dI/dS provide a measure of how many deleterious mutations will eventually be purged, given enough time. Thus, these two methods are not only independent but also provide complimentary comparisons encompassing both very strongly and weakly deleterious mutations.

We apply these approaches to large whole genome datasets from six diverse bacterial species; *Escherichia coli*, *Staphylococcus aureus, Salmonella enterica*, *Streptococcus pneumoniae*, *Klebsiella pneumoniae*, and *Mycobacterium tuberculosis*. We first draw the broad conclusion from the PSM and dI/dS analyses that the frequency of both strongly and weakly deleterious mutations in intergenic sites is intermediate between the equivalent frequencies in synonymous and non-synonymous sites. We then go on to compare selective constraint operating on different regulatory elements and examine the impact of divergence time and distance from gene borders. Strikingly, we note that selective constraint (both strong and weak) operates on an average of 69% of intergenic sites, even when the major regulatory elements are excluded.

*M. tuberculosis* is an exceptional species, in which there is little evidence of selective constraint operating on IGRs. However we demonstrate strong evidence of *positive* selection in predicted promoters of *M. tuberculosis* and posit that variation in promoters represents a key adaptive mechanism in this highly monomorphic species. We conclude that, for most species, the majority of intergenic sites are under purifying selection and moreover that mutations at these sites are commonly strongly deleterious.

## Results

### Species and data selection

We used existing large whole genome sequence datasets for six diverse bacterial species *Escherichia coli*, *Staphylococcus aureus, Salmonella enterica*, *Streptococcus pneumoniae*, *Klebsiella pneumoniae*, and *Mycobacterium tuberculosis*. Table 1 summarises the key features of the datasets and the reference genomes used in this analysis for each species. These species are diverse in terms of phylogeny (representing Gram-positive and Gram-negative taxa), in terms of population structure (ranging from the highly clonal *M. tuberculosis* to the freely recombining *S. pneumoniae*), and in terms of ecology. The *K. pneumoniae* and *E. coli* data include isolates from environmental sources and disease, the *S. aureus* and *S. pneumoniae* data includes isolates from asymptomatic carriage, and *M. tuberculosis* is an intracellular pathogen. The GC contents of these species range from 32.9% (*S. aureus*) to 65.6% (*M. tuberculosis*) (Table 1). In cases where very large datasets (1000s of genomes) were available, we sub-sampled representative strains as described in Methods. For each species, we then mapped the reads to a single reference genome (Table 1), and identified a core set of genes and IGRs; regions were included if > 90% of the sequence was present in > 95% of strains. The percentage of genes and (IGRs) included ranged from 71% (63%) in *S. pneumoniae* to 94% (93%) in *M. tuberculosis*; mean 82% (75%) (Table 1). This core genome was used in all subsequent analyses. A complete list of all isolates used in the analysis is given in Table S1.

**Table 1:** Summary of the datasets used in the analysis.

| Species | Number of isolates | Gram stain | % GC content | Reference genome | Core genes | Core IGRs | % total genes | % total IGRs | Data source |
| --- | --- | --- | --- | --- | --- | --- | --- | --- | --- |
| *E. coli* | 157 | Gram - | 50.8 | MG1655 | 4305 | 3660 | 73.50 | 64.02 | NCBI complete genomes |
| *S. enterica* | 366 | Gram - | 52.2 | Typhimurium_D23580 | 4554 | 3794 | 79.42 | 74.80 | Public Health England |
| *K. pneumoniae* | 209 | Gram - | 57.7 | NTUH_K2044 | 4787 | 4023 | 82.58 | 78.35 | Holt et al, 2015 |
| *S. aureus* | 133 | Gram + | 32.9 | HO_5096_0412 | 2405 | 2131 | 88.86 | 78.70 | Reuter et al, 2015 |
| *S. pneumoniae* | 264 | Gram + | 39.5 | ATCC_700669 | 2183 | 1864 | 71.19 | 63.09 | Chewapreecha et al, 2014 |
| *M. tuberculosis* | 144 | Gram NA | 65.6 | H37Rv | 4069 | 3157 | 93.54 | 93.44 | Casali et al, 2014 |

### Sequence properties of genes and IGRs

IGRs were identified based on reference genome annotation as described in Methods. The size distribution and GC content of both genes and assigned IGRs are shown in Figure 1. As expected, genes are significantly larger than IGRs in all species (Figure 1a). IGRs with a predicted promoter at each end tend to be larger than double terminator regions and co-oriented regions (many co-oriented regions are small spacers within operons). The GC content of IGRs is substantially lower than in protein coding sequences (Figure 1b). Although this difference is far less marked in *M. tuberculosis*, it is statistically significant in all species (p < 10^−16^, Mann-Whitney *U* test). Muto and Osawa showed that fourfold degenerate sites exhibit the widest range of GC content, whereas non-degenerate second codon positions exhibit the narrowest range; this can be explained by constraint on non-synonymous mutations at second codon positions ^22,23^. We tested for this effect by analysing synonymous, non-synonymous, and intergenic sites, and comparing the GC contents of these sites to the genome GC content for each species (Figure 1c). Our data are consistent with that of Muto and Osawa; the synonymous sites exhibit the widest range of GC content (steepest slope), and the non-synonymous sites exhibit the narrowest range of GC content (shallowest slope). The slope for intergenic sites is intermediate in gradient, indicating that constraint on intergenic sites is intermediate between that on synonymous and non-synonymous sites.

**Figure 1:**
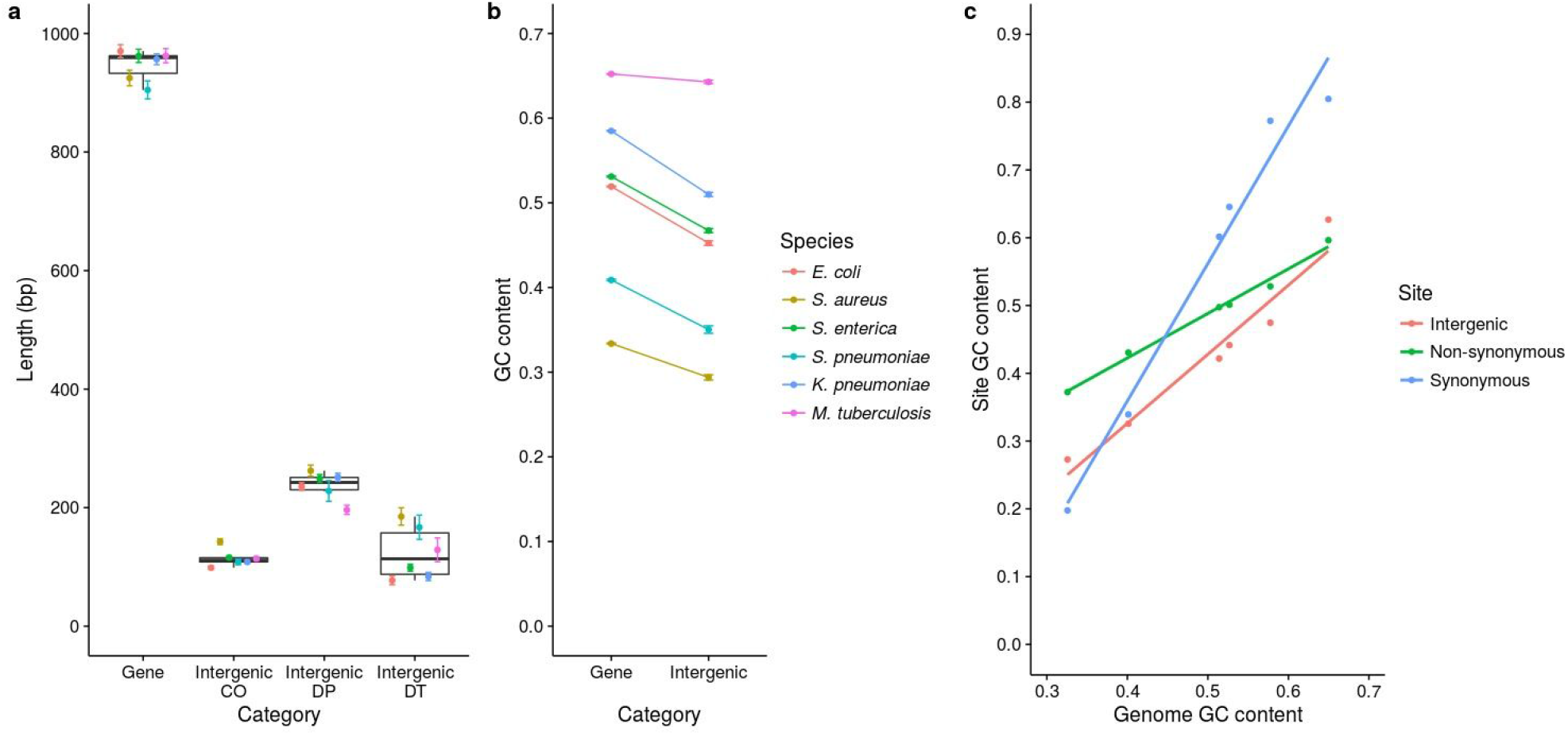
Summary of the sequence properties of genes and IGRs. Length distributions of genes and IGRs (a). IGRs were divided into three groups according to the orientation of the flanking genes: co-oriented (CO) regions are flanked by genes in the same orientation, double promoter (DP) regions are flanked by 5’ gene starts, and double terminator (DT) regions are flanked by 3’ gene ends. The points and error bars represent mean ± se. GC contents of genes and IGRs (b). GC contents were calculated for each gene and IGR individually. The points and error bars represent the mean ± se. GC contents of different site classes compared to genome GC content (c). The GC content of synonymous, non-synonymous, and intergenic sites was calculated, and compared with the genome GC content for each species. The gradient of the slope indicates the amount of constraint on the GC content of the site class (shallower slopes indicate stronger constraint).

### The Proportion of Singleton Mutations (PSM) reflects strong constraint on intergenic sites

In order to examine the frequency of strongly deleterious intergenic mutations, relative to synonymous, non-synonymous and nonsense mutations within coding regions, we first used a simple method based on site frequency spectra. Similar methods have been used on non-coding DNA in eukaryotes ^24^, and on a much smaller scale in bacteria ^5^. Mutations affecting sites under strong selective constraint are more likely to be quickly purged by selection before they begin to rise in frequency, thus are also more likely to be very rare. Here, we defined *very rare* mutations simply as those observed only once in the dataset (singletons). It is thus possible to gauge the proportion of strongly deleterious SNPs for a given site category simply by computing the proportion of those SNPs that are singletons (Proportion of Singleton Mutations; PSM). We first considered four mutation categories, intergenic, synonymous, non-synonymous, and nonsense. The mean PSM values for each of these mutation types, for all six species, are shown in Figure 2, and an analysis of all individual genes and IGRs is given in Figure S1.

**Figure 2:**
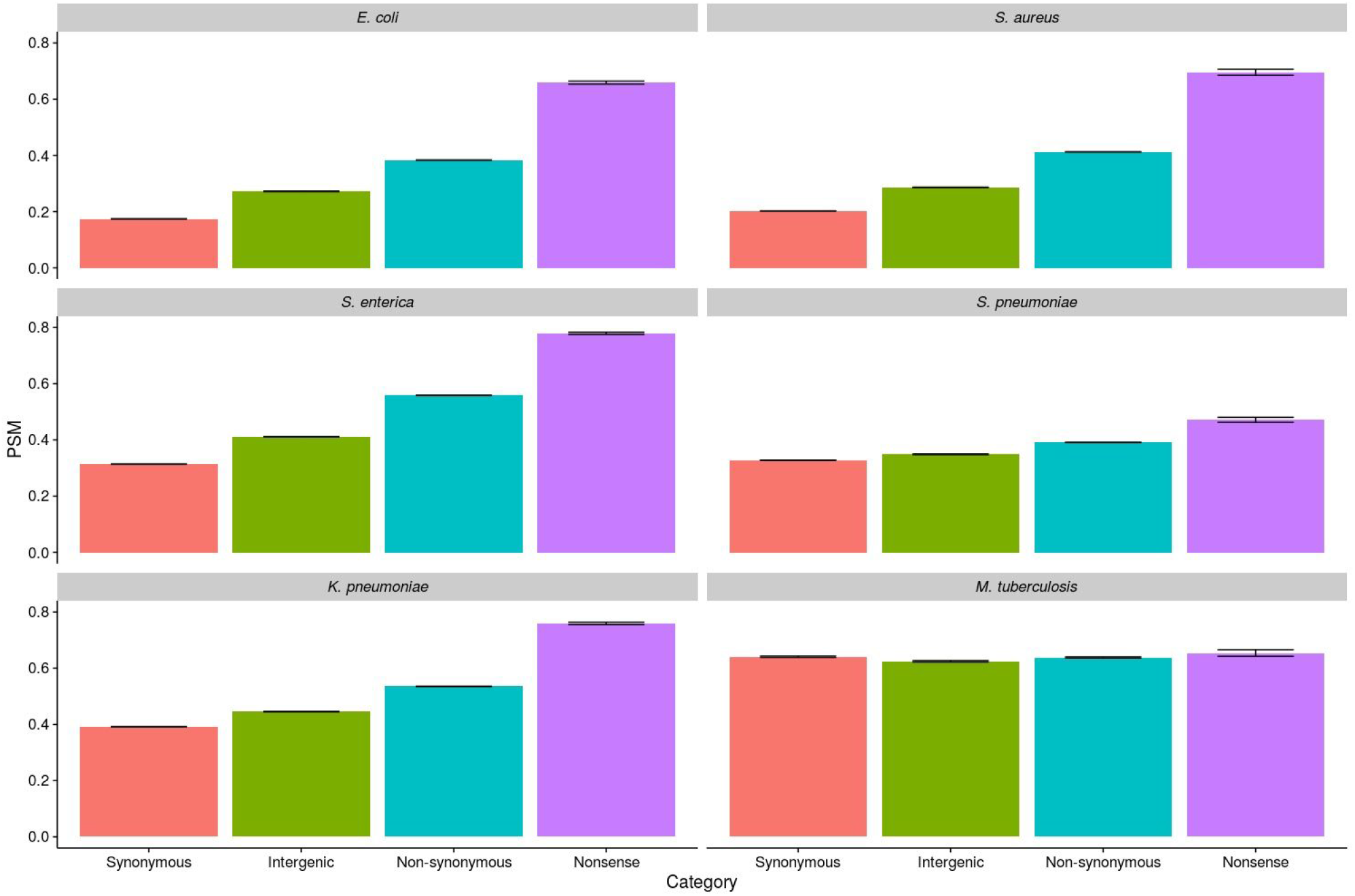
PSM (Proportion of Singleton Mutations) analysis of selection on different mutation categories. PSM values were calculated by dividing the number of singleton SNPs by the total number of SNPs within that mutation category. The error bars represent the mean ± se based on resampling the data 1000 times.

This analysis reveals a consistent trend across five of the six species, the exception being *M. tuberculosis*. Nonsense mutations had the highest PSM values, indicating the highest proportion of strongly deleterious mutations, followed by non-synonymous mutations, intergenic mutations, and finally synonymous mutations. Thus, for five of the six species, PSM values for intergenic sites were intermediate between the synonymous and non-synonymous PSM values. It follows that the proportion of SNPs at intergenic sites that are highly deleterious is intermediate between the equivalent proportions for synonymous and non-synonymous sites. Using synonymous SNPs as a baseline, the excess of highly deleterious singleton SNPs in intergenic sites, given by (PSM_I_ – PSM_S_), was on average 39% of the excess singleton SNPs in non-synonymous sites, (PSM_N_ – PSM_S_), with the range being from 31% (*S. pneumoniae*) to 46% (*E. coli*).

Although approaches based on singleton SNPs are potentially vulnerable to poor quality data, the consistency of this trend across five species means that it is highly unlikely to reflect sequencing errors. Moreover, Figure S1 shows the same trend when the data is broken down into individual genes and IGRs; therefore this observation does not reflect the presence of a relatively small number of highly conserved and unrepresentative IGRs. In *M. tuberculosis*, the exceptional species, there was very little difference between all mutation categories, and PSM scores were very high in all cases. This is consistent with very weak purifying selection across the whole genome, and/or rapid demographic expansion. Support for the former has previously been presented in an analysis of the strength of codon bias in a wide range of bacteria ^21^.

### Comparisons of dI/dS and dN/dS reveal that purifying selection on intergenic sites is time dependent

To further examine selective constraint on intergenic sites over longer time scales we extended the logic of dN/dS by computing dI/dS, where dI = intergenic SNPs per intergenic site. dI has previously been used in *M. tuberculosis* as a neutral reference ^2^. However, rather than calculate dI/dS for individual IGRs (using neighbouring genes as a source of synonymous sites), we drew pairwise comparisons by pooling sites across the whole genome, and used the genome-wide dI as the numerator and the genome-wide dS as the denominator. For each pairwise comparison we also computed genome-wide dN/dS in the same way, so as to enable comparisons between the strength of selection on intergenic sites and non-synonymous sites, relative to synonymous sites.

Previous work has shown that dN/dS values decrease with divergence time due to a lag in purifying selection, which operates much more strongly on non-synonymous than synonymous sites ^20^. We tested for the same time dependence in dI/dS by comparing pairs of very closely related genomes (within “clonal complexes” (CCs); where dS < 0.001) with those representing more distantly related genomes (“between-CCs”; dS > 0.001, Figure 3). All genome comparisons within *M. tuberculosis* were defined as “within-CC” due to the very low level of variation in this species. We also plotted, for each pair of isolates for each species, dN/dS and dI/dS against dS in order to further explore the impact of divergence time on dI/dS (Figure S2).

**Figure 3:**
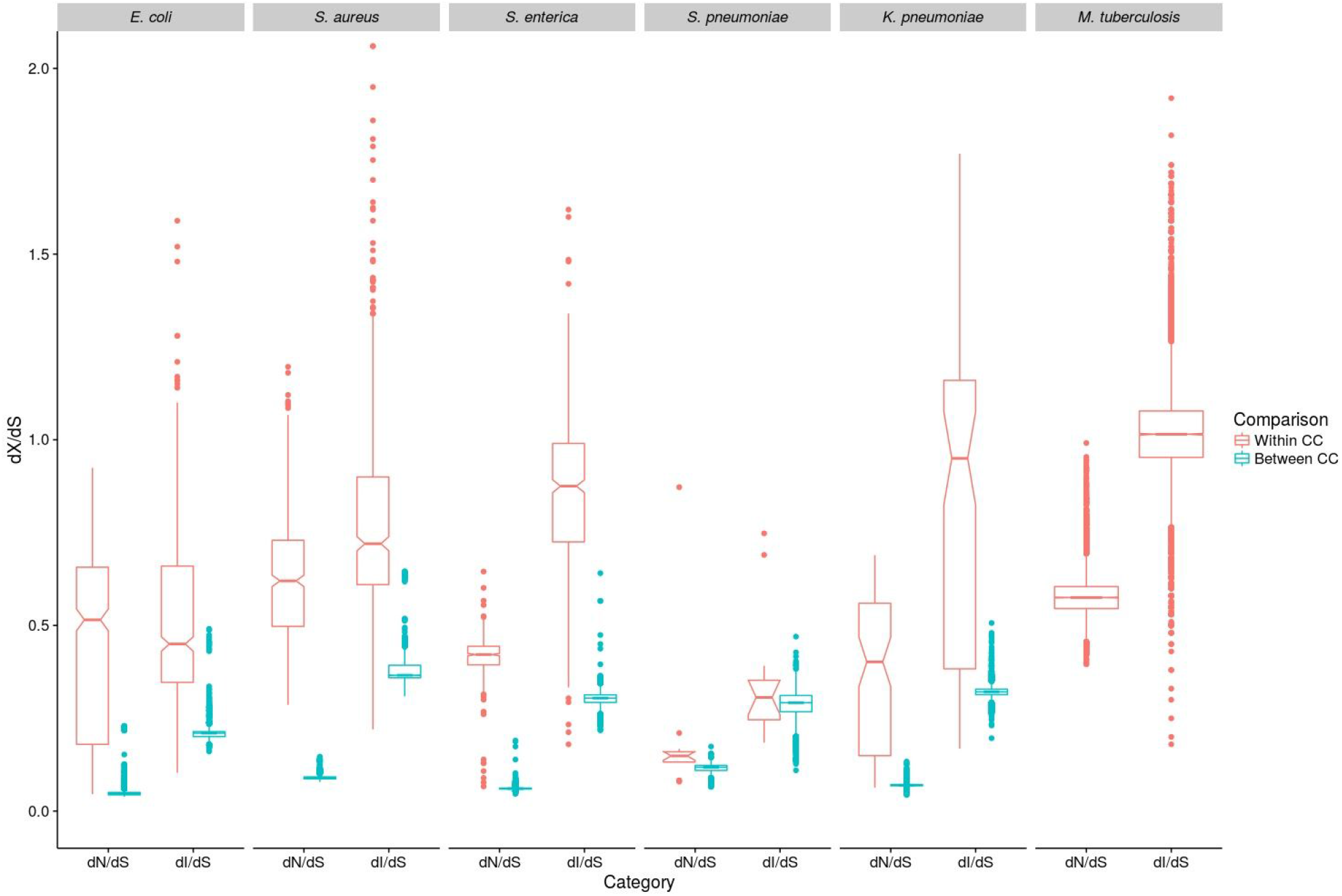
dN/dS and dI/dS analysis of selection. dN/dS and dI/dS were calculated between isolates in a pairwise manner. The results were categorised into within clonal complex (within-CC, dS < 0.001, red), and between clonal complex (between-CC, dS > 0.001, blue) comparisons, to account for the effect of divergence time on the observed levels of selection. The notches in the box plots represent 95% confidence intervals around the median. All comparisons between *M. tuberculosis* isolates were classified as “within-CC” due to the extremely low level of diversity in this species.

Figure 3 shows that, for each species, dI/dS is consistently greater than dN/dS for both within and between-CC comparisons, with the exception of within-CC comparisons for *E. coli* (between-CC comparisons for *M. tuberculosis* were not possible to compute). This confirms that the strength of selective constraint is stronger on non-synonymous sites than on intergenic sites. However, with the exception of *M. tuberculosis* and within-CC comparisons for *S. enterica,* the mean dI/dS is also consistently < 1 (p < 0.01, Wilcoxon signed rank test). This observation has previously been interpreted as evidence for positive selection on synonymous sites, which is valid only if intergenic sites are neutral ^2^. Given the results from the PSM analysis we argue instead that it confirms greater selective constraint on intergenic sites than on synonymous sites. Moreover, we note significant evidence of time dependence as predicted under our working model of purifying selection. For four of the six species (*E. coli*, *S. aureus*, *S. enterica*, *K. pneumoniae*), dN/dS and dI/dS were both significantly higher for within-CC comparisons than between-CC comparisons (p < 10^−16^, Mann-Whitney *U* test). These differences are expected if non-synonymous and intergenic SNPs are preferentially purged, relative to synonymous SNPs, over divergence time. In *S. pneumoniae*, dN/dS was significantly higher for within-CC comparisons compared to between-CC comparisons (p < 0.001, Mann-Whitney *U* test) but dI/dS was not (p = 0.82). It is possible that the signal of time dependence is weaker in this species owing to high rates of recombination. As noted above, the genetic diversity within *M. tuberculosis* is so low that between-CC comparisons were not possible, as the dS for all pairwise comparisons was < 0.001. Values of dN/dS are high within this species (mean > 0.5) and comparable to the within-CC values for *E. coli* and *S. aureus,* and the mean dI/dS value is approximately 1; these observations are again consistent with negligible purifying selection in this species.

We also plotted, for each pair of isolates for each species, dN/dS and dI/dS against dS (Figure S2). In the case of *E. coli*, *S. aureus*, *S. enterica* and *K. pneumoniae* a large number of points are evident at very low values of dS; these reflect the presence of clusters of closely related genomes in these species (i.e. clonal complexes). The absence of significant clonal clustering in *S. pneumoniae* reflects high rates of recombination, and can help to explain the lack of significant difference within and between clonal complexes in this species as noted above. However, for all species except *S. enterica* and *M. tuberculosis* there is a significant decrease of both dN/dS and dI/dS against dS (p < 10^−16^, Spearman’s correlation).

From these estimates of dI/dS and dN/dS, it is possible to calculate the number of intergenic and non-synonymous sites that are under selective constraint, relative to synonymous sites. This is derived simply as (dS-dI)/dS multiplied by the number of intergenic sites or, equivalently, (dS-dN)/dS multiplied by the number of non-synonymous sites. This is similar logic to that used previously to estimate the proportion of purged deleterious mutations in sexual eukaryotes ^25^. For this analysis, we did not discriminate between pairwise comparisons according to divergence time, all comparisons were pooled, except that we excluded *M. tuberculosis*. The species with the lowest dI/dS is *E. coli*, equating to the highest proportion of constrained intergenic sites (78.7%) whereas the highest dI/dS is in *S. aureus*; 61.0% of the intergenic sites are under selective constraint in this species. The equivalent figures for non-synonymous sites in *E. coli* and *S. aureus* are 94.6% and 89.9% respectively. Figure 4 and Table 2 give the total numbers and proportions of constrained sites in each of the five species respectively.

**Figure 4:**
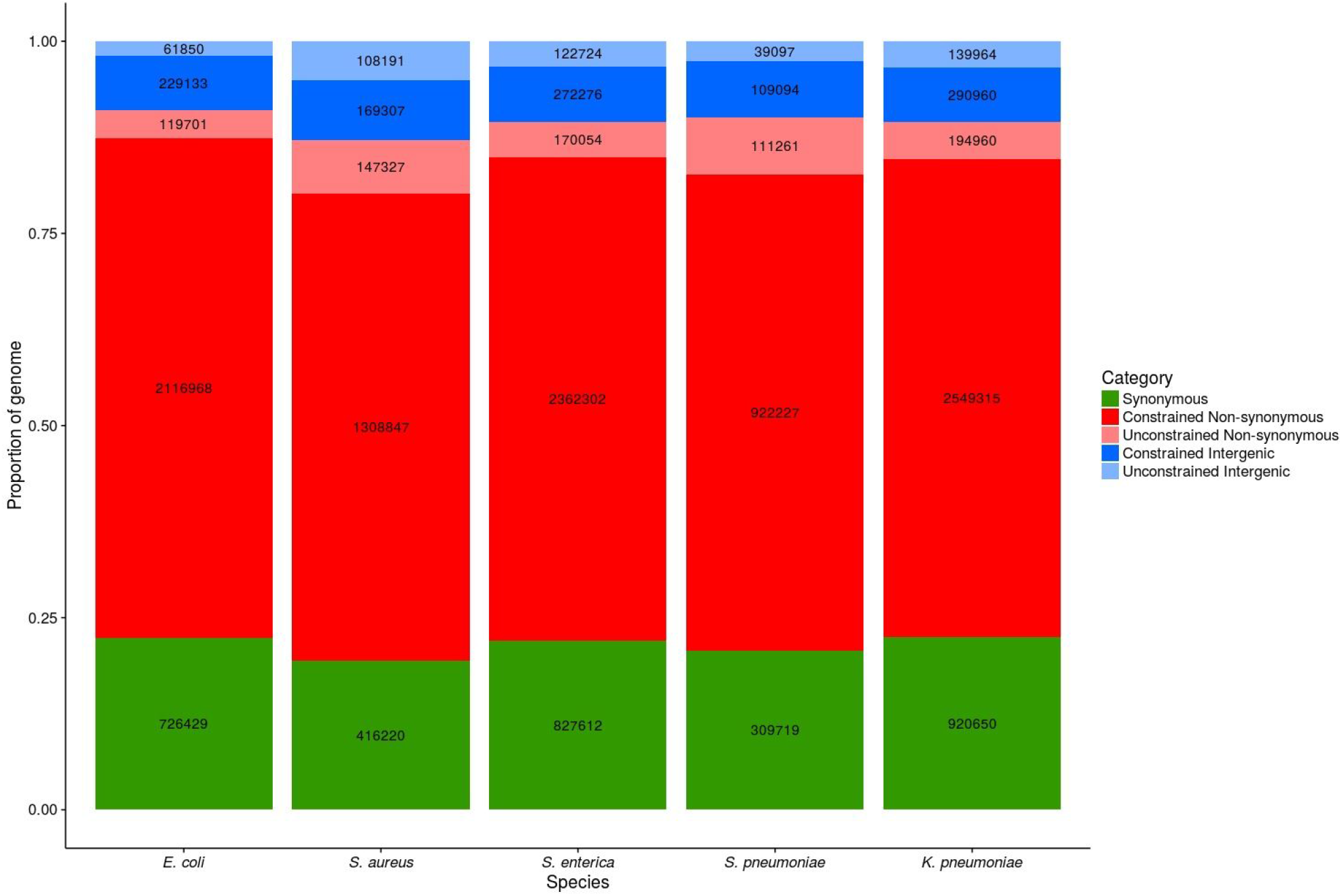
The proportions and absolute numbers of constrained and unconstrained non-synonymous and intergenic sites in the genomes of five species. The number of synonymous sites is also included to ensure that the proportion of each site is representative of the genome as a whole.

**Table 2:** Proportions of constrained sites in different genomic regions.

| Species | Non-synonymous | All intergenic | Non-coding RNA | Promoter | Terminator | Unannotated |
| --- | --- | --- | --- | --- | --- | --- |
| *E. coli* | 0.946 | 0.787 | 0.918 | 0.810 | 0.776 | 0.783 |
| *S. aureus* | 0.899 | 0.610 | 0.906 | 0.775 | 0.658 | 0.595 |
| *S. enterica* | 0.933 | 0.689 | 0.760 | 0.773 | 0.651 | 0.686 |
| *S. pneumoniae* | 0.892 | 0.736 | 0.794 | 0.851 | 0.730 | 0.732 |
| *K. pneumoniae* | 0.929 | 0.675 | 0.919 | 0.728 | 0.679 | 0.665 |
| Mean | 0.920 | 0.700 | 0.859 | 0.787 | 0.699 | 0.692 |

### Purifying selection on intergenic sites is strongest near gene starts

Although values of dI/dS < 1 are consistent with stronger selective constraint on intergenic sites than on synonymous sites, this could also arise due to slower mutation rates within IGRs than in coding regions. This might be expected if a non-negligible fraction of mutations arose during transcription, which would primarily impact on those intergenic sites nearest to the gene border ^26^. The demonstration of the time dependence of dI/dS acts to mitigate these concerns, but as a further check we calculated dI/dS values from intergenic sites immediately upstream of genes (30 bases upstream from the start codon). If intergenic sites immediately upstream of genes are transcribed, and transcription-derived mutation is a significant factor, then dI/dS should approach 1 in these regions. However, for each species (except *M. tuberculosis*) we noted the opposite; dI/dS immediately upstream of genes was in fact lower than dI/dS for intergenic sites in general (p < 10^−16^, Mann-Whitney *U* test), suggesting that transcription-derived mutation is not confounding our analysis (Figure S3). This suggests that intergenic sites close to the start of genes are under even stronger purifying selection than intergenic sites in general, which may be due to the presence of regulatory elements upstream of genes, or selection for mRNA stability to enable efficient translation ^1^. This observation is discussed further below.

### The strength of purifying selection on different classes of intergenic regulatory element

Above we demonstrate that the majority of intergenic sites, in the majority of species, are under strong selective constraint. However, we have not yet considered to what extent the strength of purifying selection may vary within a given IGR according to the presence or absence of different regulatory elements. It would be expected that sites associated with known regulatory elements should be under stronger selective constraint than sites with no known function, and it may be the case that certain classes of regulatory element are under stronger selective constraint than others. To test these expectations, we identified all non-coding RNAs, predicted promoters, and rho-independent terminators for each species (see Methods). We then applied both methods (PSM and dI/dS) to compare the strength of selective constraint on these different elements, as well as on unannotated intergenic sites.

Considering the PSM analysis (Figure S4), we note that in three species, *E. coli*, *S. aureus*, *K. pneumoniae*, SNPs occurring in non-coding RNAs were more likely to be singletons than SNPs occurring in terminators, predicted promoters or un-annotated sites. This suggests highly deleterious mutations are more common in non-coding RNAs than in other intergenic sites in these three species. However, subdividing the data in this way reduces the power of the method, and no other trends can be readily discerned. We next drew the same comparisons using dI/dS values (Figure 5). This confirmed the observation from the PSM analysis of particularly strong purifying selection on non-coding RNAs for *E. coli*, *S. aureus*, *K. pneumoniae*. Indeed, the dI/dS values for non-coding RNAs in these species are similar to the dN/dS values, suggesting that the strength of purifying selection on non-coding RNAs is similar to that operating on non-synonymous sites (Table 2, Figure 5, S4, S5). This analysis also reveals that predicted promoters are under similar levels of selective constraint to non-coding RNAs in *S. enterica* and *S. pneumoniae*, but predicted promoters are under stronger selective constraint than terminators in *S. aureus*, *S. enterica, S. pneumoniae,* and *K. pneumoniae* (p < 10^−16^, Mann-Whitney *U* test) ((Table 2, Figure 5, S5). Surprisingly, the dI/dS values for terminators are very similar to those for unannotated sites in all species. Importantly, however, it is clear that (with the exception of *M. tuberculosis*) dI/dS is < 1 in all cases, including unannotated sites, which suggests a high level of cryptic functionality.

**Figure 5:**
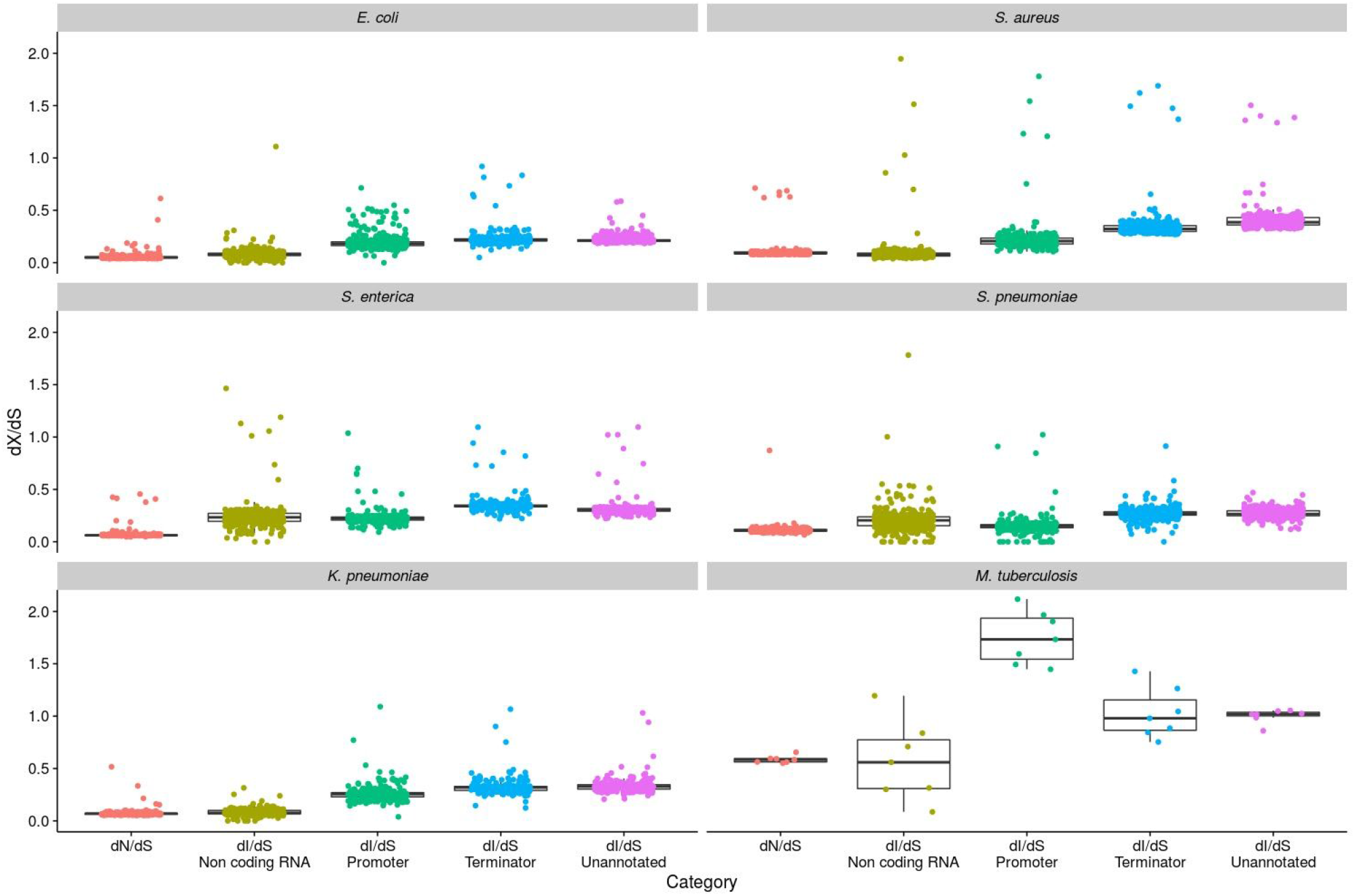
dI/dS analysis of selection on different regulatory elements. dI/dS was calculated between isolates in a pairwise manner, and the results were binned by dS (bin width = 0.0001) to control for population structure. The genome-wide dN/dS values are included to enable comparisons to be made between non-synonymous sites and the different regulatory intergenic sites.

We then computed the proportion of intergenic sites subject to selective constraint for each intergenic regulatory element, excluding *M. tuberculosis* (Table 2, Figure S5). This revealed that, on average over five species, 70% of all intergenic sites are under selective constraint. This can be broken down as follows: an average 85.9% of sites within non-coding RNAs are under selective constraint, with the equivalent figures for predicted promoters, terminators and unannotated sites being 78.7%, 69.9% and 69.2% respectively. Thus, even excluding all known regulatory elements, an average of 69% of intergenic sites in bacteria are under selective constraint. Overall, *E. coli* shows the strongest selective constraint in both intergenic and non-synonymous sites, possibly reflecting a very large effective population size; a remarkable 78.3% of the unannotated intergenic sites in this species are under selective constraint, relative to synonymous sites. These results are summarised as follows: i) non-coding RNAs tend are under very strong selective constraint (both by the PSM and dI/dS analysis) in *E. coli*, *S. aureus*, *K. pneumoniae*, ii) promoters are under stronger selective constraint than rho-independent terminators, iii) selective constraint (relative to synonymous sites) is evident in IGRs even when all known regulatory elements are excluded.

As the vast majority of intergenic sites do not correspond to any annotated features, estimates of dI/dS for the unannotated regions are similar to those when all intergenic sites are considered (Table 2, Figure S5). As discussed, this points to considerable selective constraint (relative to synonymous sites) even on intergenic sites with no known function. We noted earlier (Figure S3) that the strength of selective constraint appears to be particularly high within 30-bp of the gene boundaries. In order to further characterise the strength of constraint operating on intergenic sites for which there is no known function, we investigated SNP density in unannotated sites as a function of the distance from gene boundaries in co-oriented IGRs (where the genes flanking these regions are in the same orientation). In each species (except *M. tuberculosis*), SNP densities increase with distance from both the 5’ and 3’ gene ends (Figure 6), suggesting that the considerable cryptic functionality within these unannotated IGRs is enriched near to gene boundaries.

**Figure 6:**
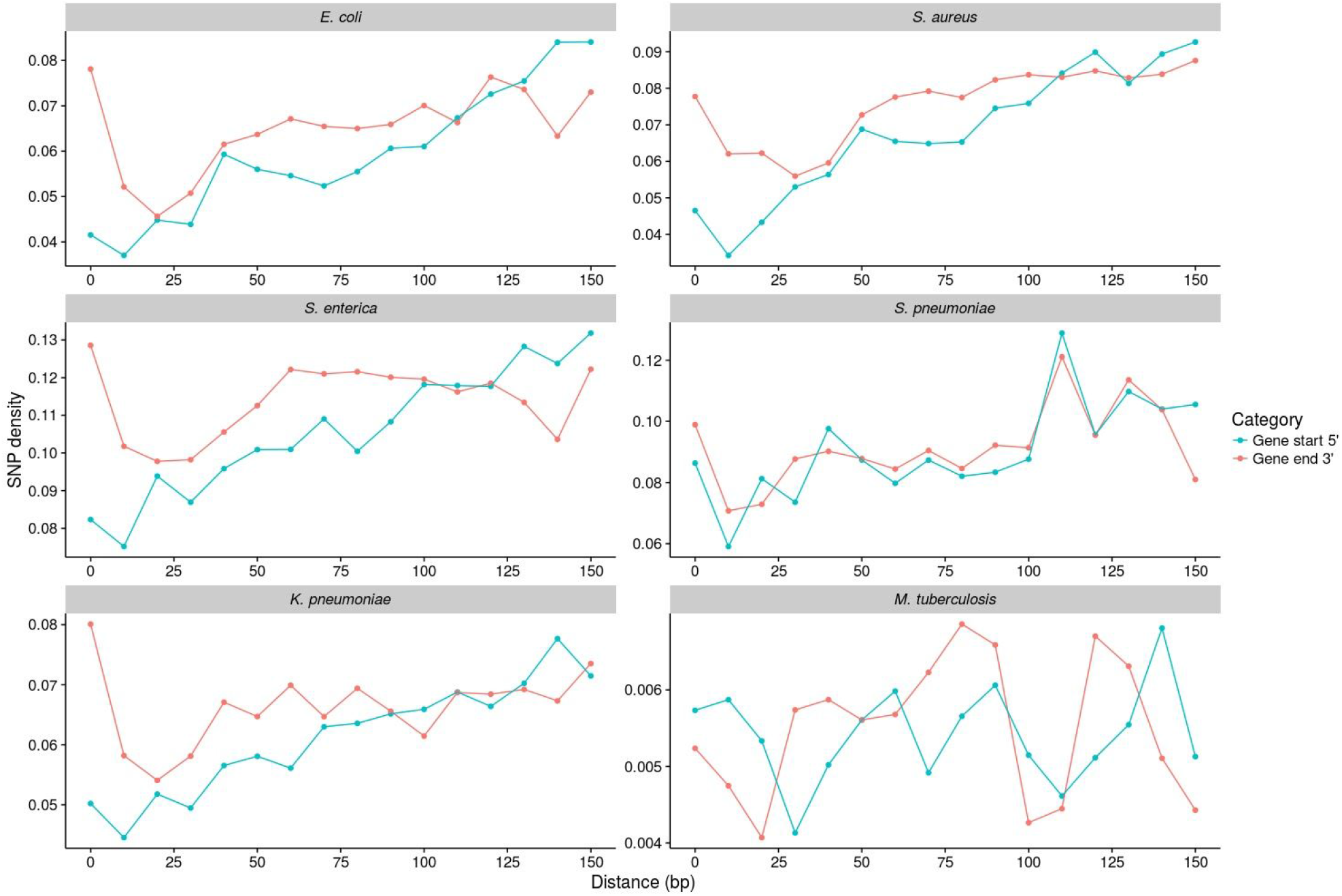
Analysis of SNP densities within co-oriented IGRs (those flanked by genes in the same orientation as each other). SNP densities were calculated in 10 bp windows by dividing the number of SNPs by the number of IGRs of that length or greater (to normalise for the unequal lengths of IGRs). Only unannotated intergenic sites were considered in the analysis.

Throughout this analysis, *M. tuberculosis* has repeatedly proved the exception and shows very little evidence of purifying selection. In fact, using the methodology presented above, we estimate that only approximately 36% of the non-synonymous sites are under selective constraint in this species, relative to synonymous sites. When all intergenic sites are considered, there is little evidence for any purifying selection using either method. However, when different intergenic regulatory elements are considered separately there is striking evidence for *positive* selection in predicted promoter regions. The mean dI/dS for *M. tuberculosis* promoters was 1.9 (Figure 5), and the distribution of dI/dS values for pairwise comparisons between isolates reveals a median of approximately 2, and that the vast majority (85%) of comparisons exhibit a dI/dS of > 1 (Figure S6). The evidence for positive selection in predicted promoter regions of *M. tuberculosis* is highly statistically significant. Of the 10513 promoter sites, 99 have experienced a SNP, compared with 6330 of the 954745 synonymous sites (p < 0.0001 by a Fisher’s exact test). We confirmed significance by resampling the predicted promoter and synonymous sites 1000 times and comparing the distributions with a z-test (p < 10^−16^).

In order to further investigate the potential functional relevance of promoter SNPs in the *M. tuberculosis* dataset, we identified genes downstream of predicted promoters harbouring SNPs (Table S2). The 71 promoter SNPs identified corresponded to 58 genes; 11 genes were identified for which the corresponding promoter harboured 2 SNPs, and one gene where the promoter harboured 3 SNPs. Many of the downstream genes are known to play a key role in virulence, resistance or global regulation. For example, 8 genes were transcriptional regulators, and promoters in four of these experienced two independent SNPs: MT0026 (a putative HTH type regulator); *CmtR* (a cadmium sensing repressor ^27^), *WhiB2* and *WhiB4* (transcription factors ^28–30^). Six genes were members of the PE/PPE protein family that are recognised virulence factors ^31^. Genes known to play a role in resistance are also identified, including one mutation in the *ethA* promoter; mutations in this promoter have previously been implicated in resistance to ethionamide ^32^. The promoter for the alanine dehydrogenase gene *ald* is the only example harbouring 3 independent SNPs, loss of function of this gene has recently been shown to confer resistance to D-cycloserine ^33^. Other notable genes include *ctpJ*, which encodes an ATPase that controls cytoplasmic metal levels ^34^, and *psk2* which plays a critical role in the synthesis of cell wall lipids ^35^. In addition, 15 hypothetical genes residing downstream of mutated promoters were identified, and in five of these cases the promoter experience two independent SNPs.

## Discussion

Here we demonstrate that intergenic site variation in bacteria is not neutral, but that on average at least 69% of intergenic sites are under selective constraint, even when all known regulatory elements are excluded. The view that IGRs mostly represent convenient neutral markers that might be used for estimating neutral mutation rates or profiles ^2^ should now be recognised as outdated and without merit; rather, these regions are rich with functional elements (many of which are unknown) and are selectively conserved and maintained. This is consistent with previous work ^11,5,6,13,14^, and also with a recent study from the Long Term Experimental Evolution project (LTEE), where a highly significant enrichment of intergenic mutations was found compared to synonymous mutations ^36^; suggesting positive selection of adaptive intergenic mutations. It is also worth noting that synonymous mutations are themselves not neutral, but are known to be under selection for various features: codon usage ^21,37^, secondary RNA structure ^1^, and possibly GC content ^23,38,39^). This means that our estimates of the proportion of sites under selective constraint are conservative, as they are computed relative to synonymous site variation.

We also find that, like dN/dS ^20,40^, dI/dS decreases with divergence time, which is evident as closely related genomes (within-CC) correspond to a higher dI/dS than more distantly related genomes (between-CC). This demonstrates that purifying selection on intergenic sites is an ongoing process over divergence time, as it is on non-synonymous sites, and that the lower prevalence of segregating sites in IGRs when compared to synonymous sites (dI/dS < 1) does not simply reflect differences in mutation rate. This is supported by the high dN/dS and dI/dS values for the within-clonal complex comparisons between isolates of *E. coli*, *S. aureus*, *S. enterica*, and *K. pneumoniae*, and by the low dI/dS values for intergenic sites immediately upstream of genes.

We also present, for the first time, a broad comparison of the strength of selection on different classes of regulatory element within IGRs. This reveals that non-coding RNAs tend to be under the strongest level of constraint, followed by predicted promoters and finally rho-independent terminators. The weaker selection observed on terminators (relative to predicted promoters and non-coding RNAs), may be a consequence of their structure. Rho-independent terminators form stem-loop structures followed by a run of uracil residues. The stability of the structure is dependent on a high level of complementarity of the two stem sequences and thus would be expected to be under strong constraint, but the loop and run of uracil residues are likely to be under weaker constraint. Our results also demonstrate that selective constraint is operating on IGRs (relative to synonymous sites) even when predicted promoters, terminators and non-coding RNAs are excluded, and that this constraint is strongest close to the 5’ and 3’ gene borders. This suggests that many functional elements in IGRs remain uncharacterised, and unannotated IGRs close to gene borders may have particular functional significance.

The power of our approaches is underscored by striking evidence for positive selection in predicted promoter regions in *M. tuberculosis*. This result is highly statistically significant, meaning that the signal of positive selection must be strong enough not to be confounded by any background purifying selection in our global comparisons. In order to gauge the functional relevance of these promoter SNPs, we identified all downstream genes, and noted a number global regulators, transcription factors, and genes implicated in virulence or resistance. A large number of hypothetical proteins were also identified, which could form targets for future studies (Table S2). This observation thus points to a key role of subtle changes within promoters for short-term adaptation through regulatory rewiring in this species, which may also help to account for the paucity of variation within coding regions.

In sum, here we have applied multiple methodologies to quantify the strength and direction of selection acting on IGRs in bacteria. We demonstrate that, on average over 5 species, at least 70% of intergenic sites are under selective constraint relative to synonymous sites, a figure which barely changes (dropping to 69%) when only considering those intergenic sites for which there is no known function. We also note the strength and direction of selection varies with the class of intergenic regulatory element, the species under consideration, distance from gene border and (possibly) the time scale of evolution; a striking case in point is highly significant evidence for positive selection within *M. tuberculosis* promoters. We conclude that our current understanding of the functions encoded in IGRs is fragmented, and we would therefore urge utmost caution before dismissing the adaptive relevance of this under-studied component of bacterial genomes. Our results call for a broad re-evaluation, and routine examination of the functional relevance of IGRs. To facilitate this, all the analysis presented in this paper can be reproduced by running a single script, using alignment and annotation files as inputs (https://github.com/harry-thorpe/Intergenic_selection_paper; full instructions available).

## Methods

### Data selection

For *S. aureus*, *S. pneumoniae*, *K. pneumoniae*, and *M. tuberculosis*, isolates were selected from recently published data ^32,41–43^. For *S. enterica*, isolates were selected from those whole genome sequenced routinely by the Gastrointestinal Bacteria Reference Unit at Public Health England. Recent large-scale bacterial genome sequencing projects have been primarily motivated by efforts to understand features of bacterial evolution which are important for public health, such as disease transmission, virulence, and antibiotic resistance. Consequently, the datasets are poorly representative of the population as a whole, with disproportionate weight given to lineages of particular clinical significance. For example the *S. aureus* data were generated as part of a retrospective study of hospital-acquired methicillin resistant S. aureus MRSA in the UK (Reuter et al. 2015), and the majority of these isolates corresponded to a single clonal lineage, CC22 (EMRSA-15). We therefore subsampled the datasets to include each major lineage and a random sample from the over-represented clonal complexes. A complete list of all isolates used in the analysis is given in Table S1.

### Sequencing, mapping and SNP calling

For each species except *E. coli*, reads were downloaded from the ENA (http://www.ebi.ac.uk/ena). For *E. coli*, completed genome sequences were downloaded from NCBI, and sheared into reads with ArtificialFastqGenerator ^44^. The isolates were mapped against a single reference genome for each species (as shown in Table 1) using SMALT (https://sourceforge.net/projects/smalt). SAMtools ^45^ was used to call and filter SNPs. Consensus fasta sequences were then used to produce an alignment for each species.

### Identification of IGRs and core genome definition

Each reference genome was annotated using Prokka ^46^. This annotation was used to extract genes and IGRs, and a core set of genes and IGRs was defined for each species (regions where > 90% of the sequence was present in > 95% of strains were included in the analysis). These genes and IGRs were used for all subsequent analysis.

### Calculation of dN/dS and dI/dS

Core gene and intergenic sequences were extracted from the alignments and concatenated to produce gene and intergenic alignments (reverse oriented genes were reverse complemented so all genes were in sense orientation). The codons within the gene alignment were shuffled, and the gene alignment was split into two (referred to as a and b). The YN00 program from the PAML suite ^47^ was used to calculate dN/dS values by the Nei and Gojobori (1986) ^48^ method in a pairwise manner for both gene alignments a and b. SNPs were counted between isolates in a pairwise manner from the intergenic alignment, and dI was calculated by dividing the number of SNPs by the length of the alignment, before applying a Jukes-Cantor distance correction ^49^. For both the gene and intergenic alignments Ns were removed from the alignment in a pairwise manner to ensure that all possible data was used. The dS values from gene alignment were used to calculate dN/dS and dI/dS, and the dS values from gene alignment b were used as a proxy for divergence time. This ensured that when plotting dN/dS and dI/dS against dS, the dS values on each axis were calculated independently, thus controlling for statistical non-independence.

### Promoter, Non-coding RNA, and terminator annotation

Promoter and terminator predictions were obtained using the PePPER webserver ^50^. Non-coding RNA annotations were obtained from the reference genome annotation GFF file produced by Prokka, where they were labelled as ‘misc_RNA’.

### Code availability

All of the code used in the analysis is available at https://github.com/harry-thorpe/Intergenic_selection_paper. The complete analysis can be reproduced by running a single script, using any alignment and annotation files as inputs. Instructions for how to do this are available in the GitHub repository.

## Acknowledgements

The *Staphylococcus aureus* genome sequences were generated as part of a study supported by a grant from the United Kingdom Clinical Research Collaboration (UKCRC) Translational Infection Research (TIR) initiative, and the Medical Research Council (Grant Number G1000803, held by Prof. Sharon Peacock) with contributions from the Biotechnology and Biological Sciences Research Council, the National Institute for Health Research on behalf of the Department of Health, and the Chief Scientist Office of the Scottish Government Health Directorate; on which EJF was a PI and SB is a postdoctoral researcher. HAT is funded by a University of Bath research studentship. The funders had no role in study design, data collection and analysis, decision to publish, or preparation of the manuscript. We are very grateful to Kathie Grant and Tim Dallman of the Gastrointestinal Bacteria Reference Unit at Public Health England for permission to use the *S. enterica* data. Finally, we thank Adam Eyre-Walker for critical reading of the manuscript and invaluable discussions.

